# *Ebony* underpins Batesian mimicry in an insect melanic polymorphism

**DOI:** 10.1101/2022.06.13.495778

**Authors:** Brodie J. Foster, Graham A. McCulloch, Yasmin Foster, Gracie C. Kroos, Jonathan M. Waters

## Abstract

The evolution of Batesian mimicry – whereby harmless species avoid predation through their resemblance to harmful species – has long intrigued biologists. In rare cases, such mimicry systems can be highly dynamic, being maintained via frequency-dependent selection on intraspecific polymorphisms, in which only some individuals within a population resemble a noxious ‘model’. Here, we use genomic approaches to identify the genetic basis of a striking mimicry polymorphism within a widespread New Zealand stonefly complex. Specifically, highly melanised specimens of *Zelandoperla* closely resemble an aposematic stonefly (*Austroperla cyrene*) well-known for its production of hydrogen cyanide. We assess convergence in the colour pattern of these two species, compare their relative palatability to predators, and use genome-wide association mapping to elucidate the genetic basis of this mimicry polymorphism. Our analysis reveals that melanised *Zelandoperla* overlap significantly with *Austroperla* in colour space, but are significantly more palatable to predators, indicating that they are indeed Batesian mimics. Analysis of 194,773 genome-wide SNPs reveals a strong outlier locus (*ebony*) differentiating melanic (mimic) versus non-melanic phenotypes. As *ebony* has a well-documented role in insect melanin biosynthesis, our findings highlight its conserved function across deeply divergent hexapod lineages. Distributional records suggest a link between the occurrence of *Zelandoperla* mimics and forested ecosystems where the model *Austroperla* is abundant, suggesting the potential for adaptive shifts in this system underpinned by environmental change.

## Introduction

Batesian mimicry systems – whereby harmless species avoid predation by resembling harmful species – have long intrigued evolutionary biologists (Bates 1862; Joron & Mallet, 1998). In particular, researchers have sought to understand the genetic mechanisms by which seemingly intricate ‘warning’ colouration patterns can evolve repeatedly across independent lineages (Ferguson *et al*. 2011; Kunte *et al*., 2014), and the potential adaptive role of intermediate phenotypes which have been thought to represent evolutionary ‘stepping stones’ (Ruxton et al., 2004; Kikuchi & Pfennig, 2010). Additionally, the rare occurrence of Batesian polymorphisms, in which only some members of a population resemble a noxious ‘model’ (Mallet & Joron, 1999; Tsurui-Sato *et al*., 2019), leads to questions regarding the potential key role of frequency-dependent selection as a driver of evolutionary diversification.

Aposematism (warning colouration) and mimicry are frequently characterised by bright colours displayed against a background of black melanic pigmentation (Ruxton *et al*., 2004). Such high-contrast patterns are thought to promote recognition and avoidance learning by predators (Aronsson & Gamberale-Stille, 2013; Zvereva & Kozlov, 2016). While genes responsible for adaptive melanin pigmentation have been well characterised in a few well-studied insect genera (Rebeiz *et al*., 2009; Nadeau *et al*., 2016; van’t Hof *et al*., 2019; Villoutreix *et al*., 2020), additional data are needed to test for conservation of their roles across more divergent insect lineages (Liu *et al*., 2016; van’t Hof *et al*., 2019). While early genetic studies of animal colouration relied heavily on candidate gene approaches (Protas & Patel, 2008; Hoekstra, 2006), more recent mapping approaches have led to the discovery of previously unknown loci associated with colour variation (Gautier *et al*., 2018; Nosil *et al*., 2018). Recent fine-scale mapping in the peppered moth, for instance, revealed that industrial melanism is caused by a transposable element insertion into the gene *cortex* (van’t Hof *et al*., 2016), rather than by mutations in well-characterised insect melanin biosynthesis genes (Wittkopp *et al*., 2002; Arakane *et al*., 2009; van’t Hof & Saccheri, 2010).

In this study, we test for the role of insect pigmentation genes in an unusual mimicry polymorphism (Fig. 1, 2*a*) recently detected in Plecoptera (stoneflies), an ancient lineage of aquatic insects (Tong *et al*., 2015; Evangelista *et al*., 2019). Specifically, black melanic pigmentation (Fig. 1) is the major component of adult colouration in a chemically defended New Zealand stonefly, *Austroperla cyrene* (Newman) (Austroperlidae) (McLellan, 1997). Similarly distinctive melanic ‘mimic’ phenotypes have been reported in two co-distributed but apparently harmless stonefly species (McLellan, 1998; McLellan, 1999). Specifically, putative mimics in the *Zelandoperla fenestrata* species complex (Gripopterygidae) are highly melanised, with conspicuous white and yellow features similar to those in *Austroperla* (Fig. 1; McLellan, 1999). Such distinctive colouration is otherwise extremely unusual among stoneflies, having been recorded in only in a handful of aposematic Austroperlidae species from Australia and South America (Foster *et al*., 2021).

**Figure 1.**
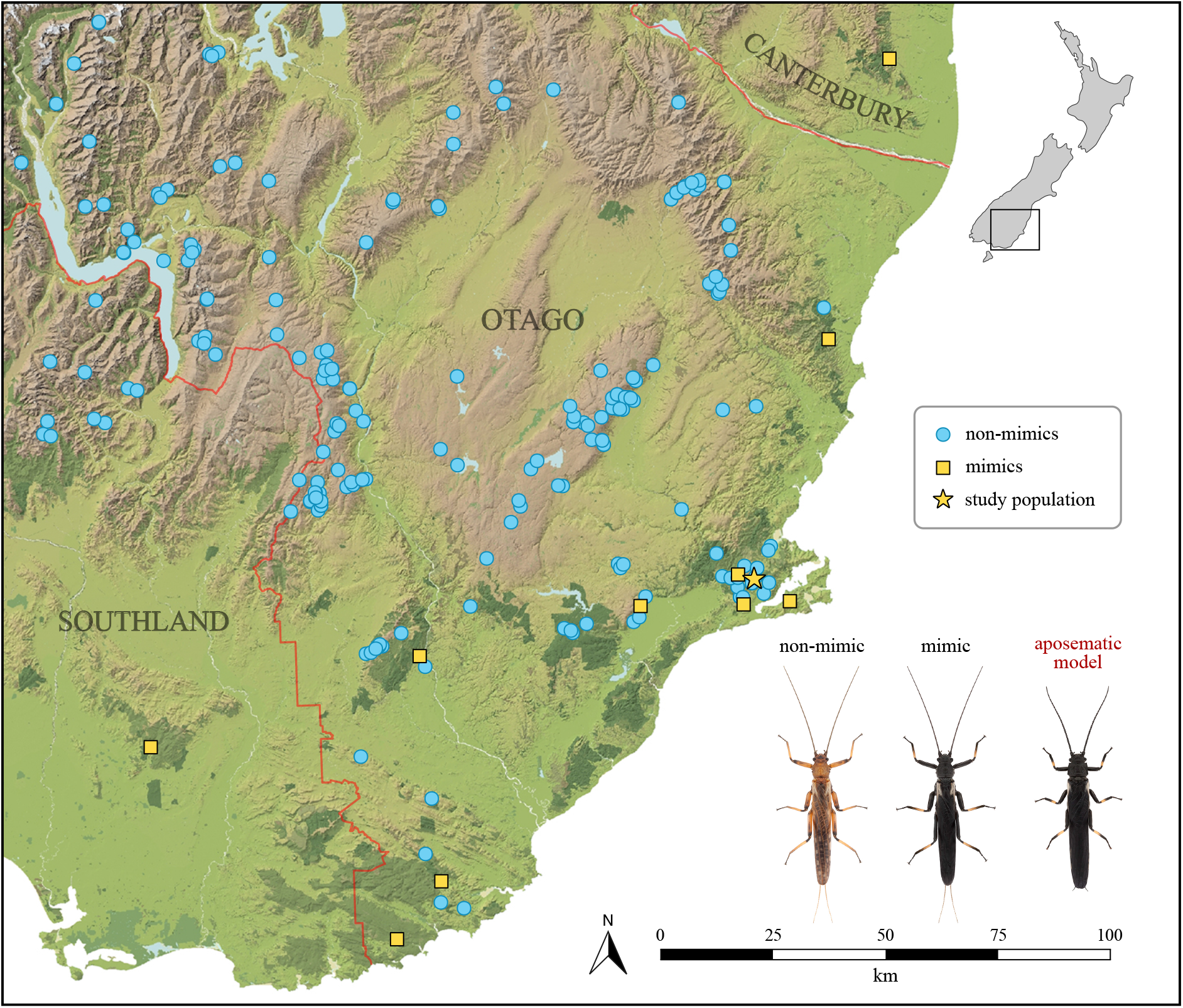
Southern New Zealand locations where mimics of *Austroperla cyrene* were identified (yellow) among adult collections of the *Zelandoperla fenestrata* species complex (blue). The location of the study population is indicated with a star. Historical distributional records suggest an association between forested habitats (dark green; where the noxious model *Austroperla* is abundant; Townsend et al., 1997; Nyström et al., 2003) and the presence of melanic *Zelandoperla* mimics.

**Figure 2.**
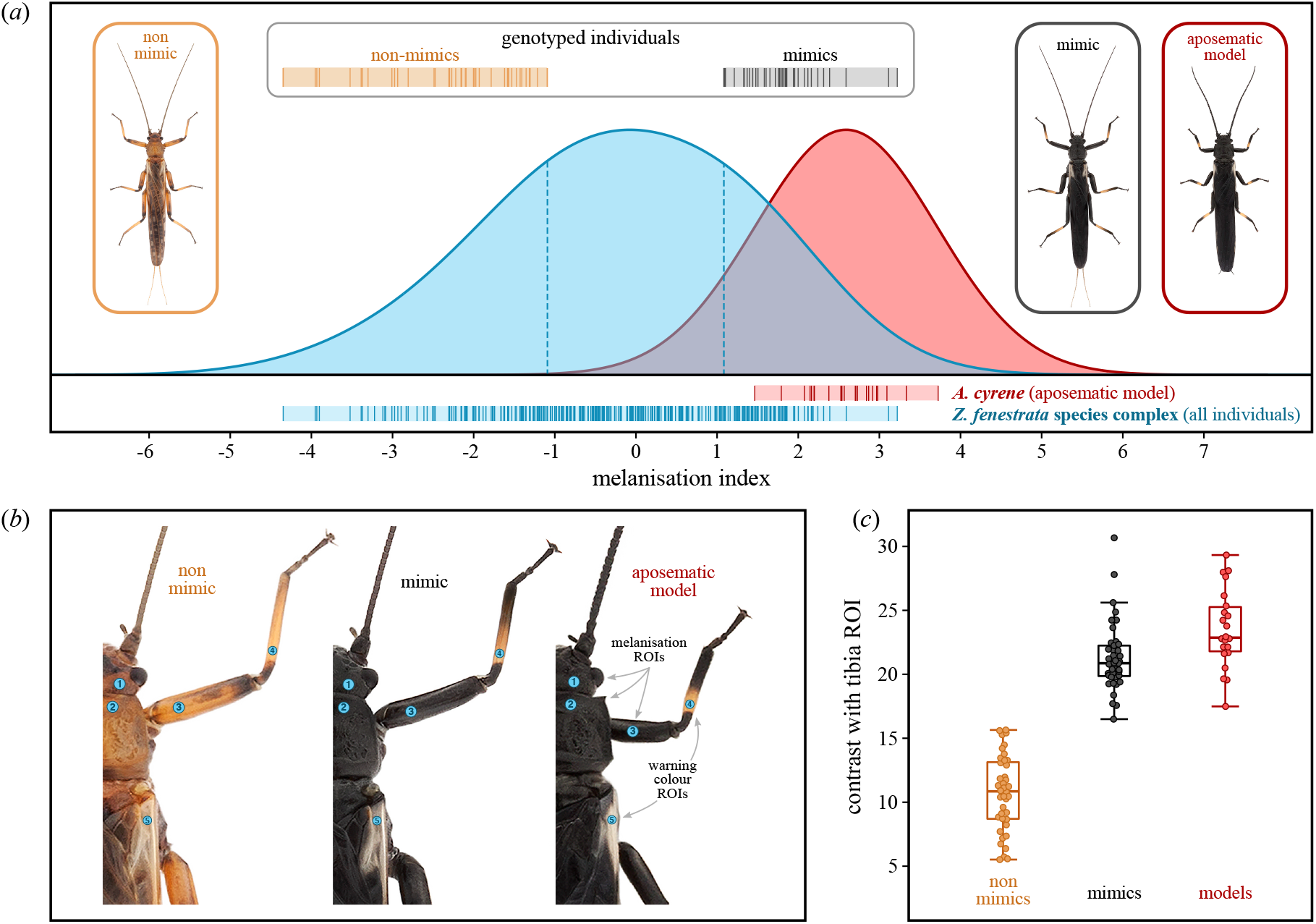
(*a*) Density estimates of overall body melanisation in a colour-polymorphic population of the *Zelandoperla fenestrata* species complex (blue) and the aposematic model (*Austroperla cyrene*; red). Individual values of melanisation are highlighted for mimics and non-mimics included in genome-wide association (GWA) analysis. (*b*) Locations and sizes of regions of interest (ROIs) sampled to obtain colour measurements, representing body melanisation (ROIs 1-3) and the brightness of warning colouration in *Austroperla* (ROIs 4 and 5). (*c*) Contrast (average differences in brightness) between body melanisation and warning colouration on the tibiae of the aposematic model compared with equivalent areas in mimics and non-mimics included in the GWA analysis.

The strong resemblance of melanic *Zelandoperla* individuals to *Austroperla* (Fig. 1), despite their ∼60 million years of divergence (McCulloch *et al*., 2016), is hypothesised to constitute Batesian mimicry (McLellan 1999). Nevertheless, the palatability of mimics remains to be experimentally demonstrated to rule out the possibility that both species are aposematic (Müllerian mimicry; Ruxton *et al*., 2004). Under current taxonomy, mimics and other dark-coloured individuals in the *Z. fenestrata* species complex have until now been recognised as *Z. tillyardi*, and light-coloured individuals as *Z. fenestrata* (McLellan, 1999). Beyond their differences in colouration, however, no substantial morphological (McLellan, 1999) or single-locus genetic (McCulloch *et al*., 2009; McCulloch, 2011) differentiation has been detected between these forms, suggesting that they may actually represent a single polymorphic species.

The melanic polymorphism in *Zelandoperla* (Fig. 1) provides an ideal system for assessing the ecological and genomic bases of mimicry. Here, we conduct detailed analyses of colour variation in *Zelandoperla* and *Austroperla* to test the hypothesis that these taxa exhibit phenotypic overlap consistent with mimicry. We estimate the prevalence of mimicry in polymorphic *Zelandoperla* and assess the importance of individual components of mimetic colouration toward resembling the model species. We also use distributional data to assess potential associations between *Zelandoperla* melanism and forested habitats where *Austroperla* is abundant (Townsend *et al*., 1997; Nyström *et al*., 2003). Furthermore, we use predation experiments to determine the type of mimicry exhibited by highly melanised individuals, and employ a genotyping-by-sequencing (GBS) approach to identify the genomic basis of this rare mimicry polymorphism.

## Materials and methods

### (a) Distribution of mimicry

*Austroperla* mimics within the *Z. fenestrata* species complex reportedly occur in Otago in the South Island of New Zealand (Tillyard, 1923) but no other information about their distribution is available. To better understand the prevalence of putative mimicry and to locate suitable populations for study, we examined the colouration of adult collections of the *Z. fenestrata* species complex (*Z. fenestrata, Z. tillyardi*, and *Z. pennulata*) from across New Zealand. A total of 1,696 ethanol-preserved and pinned specimens (810 females, 886 males) were examined across 430 unique locations originating from recent field collections, two private collections, and five institutional collections: Auckland War Memorial Museum, New Zealand Arthropod Collection (Auckland), Museum of New Zealand Te Papa Tongarewa (Wellington), Canterbury Museum (Christchurch), and Otago Museum (Dunedin) (Supplementary Fig. 1).

### (b) Sampling

Sampling focused on a population in the Water of Leith catchment of Dunedin (−45.8231, 170.5054), as both light-and dark-coloured forms of the *Z. fenestrata* species complex have been recorded in this catchment (previously ascribed to *Z. fenestrata* and *Z. tillyardi*, respectively, in McLellan, 1999). Sampling was undertaken in areas above and below the confluence of the Water of Leith and Morrisons Burn (Supplementary Fig. 2) during the peak emergence periods (November-February) of 2017-2018 and 2018-2019. Last instar nymphs of the species complex were collected from stones and wood in shallow stream riffles and maintained in the laboratory under an ambient cycle at 11 °C until they reached adulthood (McCulloch *et al*., 2019). A total of 338 adults (192 females, 146 males) were obtained for colour measurements and selection of individuals for genotyping. The aposematic model species (*Austroperla*) is also abundant at this site, and 22 reared adults (14 females, 8 males) were obtained for colour measurements to provide a reference with which to assess the strength of mimetic resemblance in individuals from the study population.

### (c) Analysis of melanisation

Live adults of the *Z. fenestrata* complex and *Austroperla* were imaged in the human visible spectrum approximately 28 hours after emergence to ensure that individuals were fully coloured. Digital photography was preferred to obtaining measurements of colour and reflectance using spectrophotometry, as photography is more practical for live animals and permits sampling of any number of regions of interest (ROIs) for analysis. Further, reflectance in the visible spectrum is known to capture variation in patterns of melanisation (Pum Lee & Wilson, 2006; Stavenga *et al*., 2012), which form the basis of mimetic resemblance between the two species. Photographs were taken in RAW format using a Canon 40D camera with an EF-S 60 mm f/2.8 lens under fixed magnification, exposure time, aperture, and ISO sensitivity settings (see Supplementary Methods), with uniform lighting provided by a Schott-Fostec DCR II 150 W fibre optic illuminator diffused through a polystyrene foam cylinder. Stoneflies were photographed at a standardised distance and angle on a 18% reflectance neutral grey background, and in each image a matte ColorGauge Pico standardised colour target (Image Science Associates, Williamson, NY, USA) was included to enable objective measurements of colour. Image files were initially normalised to the nominal reflectance of the neutral grey background using the Multispectral Image Calibration and Analysis (MICA) toolbox (Troscianko & Stevens, 2015). The average red, green, and blue (RGB) colour channel values of each image were then linearised and equalised in reference to the nominal RGB values of the grey patches of the ColorGauge Pico target in Adobe Photoshop CC 2021 (Adobe Systems, San Jose, CA, USA).

Colour measurements were taken from calibrated images in five regions of interest (ROIs) using the RGB Measure function in ImageJ (Schneider *et al*., 2012). The first three chosen ROIs correspond to areas of the head (1), pronotum (2), and femorae (3), to provide a broad representation of overall body melanisation. The additional ROIs correspond to areas on the foreleg tibiae (4) and forewing bases (5) associated with warning colouration in the aposematic model *Austroperla* (Fig. 2*b*). Mean RGB values from each ROI were converted to HSB (hue, saturation, brightness) values using the *colorsys* module in Python, and only the achromatic brightness values of the five ROIs, within a range of 0% (black) to 100% (white), were used in subsequent analyses. A principal component analysis (PCA) was performed on brightness data from the three ROIs associated with overall body melanisation in a combined dataset (*n* = 360) containing individuals from the *Z. fenestrata* complex and the aposematic model *Austroperla*. The first principal component (PC) accounted for 81.8% of the total variance in brightness (Supplementary Fig. 3), which was contributed to equally by the first three ROIs (34.0%, 33.5%, and 32.5%, respectively), indicating that scores on the first PC provide a reliable measure of overall melanisation. We subsequently used scores on the first PC as an index of melanisation for assessing the broad resemblance of individuals in the *Z. fenestrata* complex study population to those of the model species.

Individuals of the *Z. fenestrata* complex were selected for genotyping based on their position in the melanisation index. Distributions of melanisation were visualised separately for the *Z. fenestrata* complex study population and *Austroperla* using kernel density estimation in the R package *ggplot2* (Wickham, 2016). Variation in melanisation in the study population was continuously and normally distributed (Shapiro-Wilk normality test: W = 0.9833, *p* < 0.01) with many intermediates, and overlapped significantly with the variation observed in *Austroperla* (Fig. 2*a*). As GWA analyses of quantitative traits with continuous variation may lack statistical power to detect loci with small phenotypic effects, we followed the approach described by Kardos *et al*. (2016) to maximise phenotypic variance by restricting the analysis to individuals with extreme colour phenotypes. Of the 338 individuals for which colour measurements were available, we selected 48 of the most highly melanised individuals (hereafter ‘mimics’) and 48 of the least melanised individuals (hereafter ‘non-mimics’) for genotyping.

### (d) Warning colouration and palatability

The resemblance of some individuals of the *Z. fenestrata* species complex to *Austroperla* appears to be most simply captured by overall levels of body melanisation. However, unmelanised areas on the tibiae and forewings, associated with warning signals in *Austroperla*, may also constitute major components of the resemblance between the two species. To understand how these features might contribute to mimicry, we investigated how the brightness of warning colour ROIs on the tibiae and forewings relates to overall body melanisation in the *Z. fenestrata* species complex using linear regression (LR) models with Bonferroni corrections. Next, we considered the combined effects of melanism and warning colouration by comparing levels of contrast (differences in achromatic brightness) between *Austroperla* and 41 mimics and 42 non-mimics that were retained in the GWA analysis. Finally, to determine whether mimicry of *Austroperla* by individuals of the *Z. fenestrata* species complex acts in a Batesian or Müllerian manner, we conducted a palatability experiment using individuals from the study population as prey and semi-aquatic spiders (*Dolomedes aquaticus* Goyen; Pisauridae) as predators.

*Dolomedes aquaticus* is a natural predator of adult stoneflies and relies on mechanical stimuli rather than vision for prey capture (Williams, 1979), making it an ideal experimental predator for assessing prey palatability without potentially confounding effects from visual cues. Fifteen subadult and adult female *D. aquaticus* were collected from an unforested stream on the Kakanui Range (−44.9671, 170.3258) where *Austroperla* had not been detected despite extensive sampling, and as such were assumed to be naïve to the chemical defences of *Austroperla*. Spiders were acclimatised to the laboratory environment and regular feeding trials were undertaken in which spiders were randomly allocated mimics, non-mimics, or models (*Austroperla*) as prey (see Supplementary Methods for experimental procedures). Visual recognition of prey was made impossible by conducting feeding trials during the dark cycle, and the palatability of each prey type was measured as the percentage of the live mass of each prey item that was consumed by predators. Differences in the palatability of 19 mimics, 20 non-mimics, and 14 models were assessed using a linear mixed effects (LME) model implemented in the R package *lmerTest* (Kuznetsova *et al*., 2017), with a Tukey’s HSD test used to make post-hoc pairwise comparisons. To account for repeated measures of palatability by individual spiders, predator ID was included as a random effect in the LME model and its significance was assessed using a likelihood ratio test (LRT).

### (e) Genotyping data

Genomic DNA was extracted from the head and femur tissue of 96 individuals from the *Z. fenestrata* complex study population (48 mimics and 48 non-mimics) using a DNeasy kit (Qiagen) following the manufacturer’s protocols. Genotyping-by-sequencing (GBS) libraries were prepared following the protocol described by Elshire *et al*. (2011) using the restriction enzyme *Ape*KI (New England Biolabs, Ipswich, MA, United Sates). Libraries were then size-selected (243-500 bp range) using a Pippin Prep (Sage Science, Beverly, MA, United States) and sequenced on an Illumina HiSeq 2500 (Illumina, San Diego, CA, Unites States). Raw sequence reads were demultiplexed and barcodes were removed using the *process_radtags* module of Stacks v. 2.53 (Rochette *et al*., 2019). Reads were then trimmed to a common length of 72 bp and mapped to a draft genome assembly of a conspecific individual from the species complex (McCulloch *et al*., 2021) using *Burrows-Wheeler Aligner* v. 0.7.15 (*MEM*; Li & Durbin, 2009). SNPs were called using the Stacks reference-based pipeline and the *populations* module was used to output genotype data for downstream analyses. Filtering was applied in VCFtools v. 1.16 (Danecek *et al*., 2011) to retain only biallelic loci that were present in at least 50% of individuals with a read depth between 2 and 30 and a minor allele frequency ≥0.01. To assess the suitability of the population sample and complete SNP dataset for the GWA analysis, we also created a more stringently filtered ‘neutral’ dataset containing only SNPs that were in Hardy-Weinberg equilibrium and present in at least 80% of individuals.

### (f) Population genomic analyses

Principal component analysis (PCA) was used to assess whether genetic structure exists in the study population that could confound the detection of outlier loci. PCAs of the complete and neutral SNP datasets were conducted in PLINK v. 1.9 (Purcell *et al*., 2007) and visualized using the R package *ggplot2* (Wickham, 2016). Thirteen individuals (7 mimics and 6 non-mimics) identified as outliers in the PCAs were removed from subsequent analyses, and the SNP calling and filtering methods described previously were repeated for the remaining 83 samples. Next, we repeated the PCA for the neutral dataset and quantified levels of genome-wide differentiation using the R package *hierfstat* (Goudet, 2005). Pairwise F_ST_ values were calculated between mimics and non-mimics, and among three groups of individuals originating from different locations of the study area. Specifically, we aimed to determine (i) if mimics (*n* = 41) and non-mimics (*n* = 42) exhibited genetic differentiation from each other; and (ii) if adjacent populations sampled from the upper regions of the Water of Leith (*n* = 47), from Morrisons Burn (*n* = 29), and beneath the confluence of the two streams (*n* = 7; Supplementary Fig. 2), exhibited genetic differentiation from each other. Finally, we used the *boot*.*ppfst* function in *hierfstat* to obtain 95% confidence intervals from 20,000 bootstrapping replicates of the pairwise F_ST_ comparisons, and considered genetic differentiation to be statistically significant if the lower-bound confidence interval (CI) of pairwise F_ST_ estimates did not overlap with zero.

### (g) Genome-wide association scan

We searched for loci potentially associated with melanism using BayeScan v. 2.1 (Foll & Gaggiotti, 2008), an outlier detection method that uses allele frequency differences between populations (here, mimics and non-mimics) to identify individual candidate loci potentially under selection. BayeScan estimates a posterior probability (α) of a given locus being under selection by defining two alternative models (one with and one without selection), where positive values of α indicate diversifying selection and negative values indicate balancing or purifying selection (Foll & Gaggiotti, 2008). BayeScan was run using default parameters, and model convergence was confirmed based on trace plots and Geweke’s diagnostic tests (Geweke, 1992) executed in the R package *coda* (Plummer *et al*., 2006). Loci with α values significantly greater than zero and q-values less than 0.05 (corresponding to a false discovery rate of 5%) were considered to be outliers and were mapped to the reference genome assembly (McCulloch *et al*., 2021). Genes located within 50 kb of an outlier locus were identified from an existing functional annotation of the reference genome based on the eggNOG database (Huerta-Cepas *et al*., 2016) and homology with *Drosophila melanogaster* genes.

### (h) Sanger sequencing validation

Sanger sequencing was used to validate GBS genotypes observed at a BayeScan outlier locus due to its location within *ebony*, a strong candidate gene in the melanin biosynthesis pathway of insects (Wright, 1987; Wittkopp *et al*., 2002). Primers were designed using *Primer3* (Untergasser *et al*., 2012) to amplify a 265-bp fragment surrounding the outlier SNP for all 96 individuals initially included in the GBS analysis. Polymerase chain reactions (PCRs) were run in 12.5 µl volumes, each containing 5 µl water, 5 µl of MyTaq HS Red Mix (Bioline, London, UK), and 0.24 µL each of 10 mM forward (eb23532F; CTTGTTGACGTACAGCTCAAGTG) and reverse primers (eb23815R; CTTCCTTGAGTGCAGCTTGC). Cycling conditions involved an initial denaturation step (95°C for 4 minutes) followed by 35 cycles of 95°C for 30 seconds, 55°C for 30 seconds, and 72°C for 30 seconds, with a final annealing step of 72°C for 10 minutes. Amplification was verified by agarose gel electrophoresis, and samples were cleaned with 1 µl of Exonuclease I (New England Biolabs Inc, Ipswich, United States) and 1 µl of Antarctic Phosphatase (New England Biolabs Inc, Ipswich, United States). Sanger sequencing was conducted by the University of Otago Genetics Analysis Service (https://gas.otago.ac.nz/), using an ABI 3730xl DNA Analyser. Sequence chromatograms were edited and aligned in Geneious v. 11.1.5 (Kearse *et al*., 2012) using the MUSCLE plugin (Edgar, 2004), and JModeltest v. 2.1.2 (Darriba *et al*., 2012) was used to select the best model of sequence evolution under the AIC criterion. Phylogenetic relationships among *ebony* haplotypes were reconstructed in Garli v. 2.01 (Zwickl, 2006) under the T92 + G model of evolution with indels treated as missing data.

## Results

### (a) Elements of mimicry

Analysis of the colour of 360 live adult stoneflies in this study confirmed that melanism in the *Z. fenestrata* species complex overlaps significantly with levels of melanisation observed in *Austroperla* (Fig. 2*a*). Highly melanised individuals within this area of overlap were considered to be mimics of *Austroperla* and represented approximately 15% of the sampled population. Mimics were found to display significantly higher internal pattern contrast than non-mimics owing to ‘warning colour ROIs’ on the tibiae and forewings (Fig. 2*b*) remaining relatively unmelanised even in highly melanised individuals (Fig. 2*c*). Increased body melanisation was associated with significant reductions in the brightness of the forewing ROI (LR: Bonferroni adjusted *p* < 0.001; Supplementary Fig. 4*b*) but had no effect on the brightness of the tibia ROI (LR: Bonferroni adjusted *p* > 0.05; Supplementary Fig. 4*a*). As a result, the bright bands on the tibiae of mimics were found to contribute to higher contrast and greater resemblance of the model than forewing colouration (Supplementary Fig. 4*b, d*). Significant differences were found in the palatability of prey as perceived by predators in feeding experiments (LME: F = 1227, df = 2, *p* < 0.001) with no effect of repeated measures by individual spiders detected (LRT: *p* > 0.05). Spiders consumed, on average, 91.3% of the mass of mimics, 89.6% of non-mimics, and 5.6% of models (*Austroperla*) (Supplementary Fig. 5). Mimics and non-mimics did not differ in their apparent palatability (Tukey’s HSD: *p* > 0.05) and both were significantly more palatable than models (Tukey’s HSD: *p* < 0.001), indicating that the resemblance of some highly melanised *Zelandoperla* to *Austroperla* is consistent with Batesian mimicry.

### (b) Distribution of mimicry

Analysis of extensive collections of *Zelandoperla* across New Zealand revealed highly melanised ‘mimic’ individuals from 11 locations in southeast New Zealand (Fig. 1; Table S1). These melanic individuals were almost exclusively associated with densely forested lowland habitats where *Austroperla* is known to be highly abundant (Townsend *et al*., 1997; Nyström *et al*., 2003).

### (c) Genome-wide differentiation

GBS yielded a total of 255 million reads across 96 individuals, which were aligned to the reference genome at an average mapping rate of 96.3%. After filtering, 83 individuals containing 194,773 SNPs were retained in the complete dataset and 7,468 SNPs were retained in the neutral dataset. Genetic differentiation in the neutral SNP dataset was low with no evidence of population substructure (Fig. 3*a*). Additionally, we found no significant F_ST_ differentiation between mimics and non-mimics (F_ST_ = 0, lower-bound CI < 0) or between individuals sampled from different areas of the study site (F_ST_ = 0.0009, lower-bound CI < 0), indicating that the complete SNP dataset is well suited for detecting outlier loci against a homogenous genetic background.

**Figure 3.**
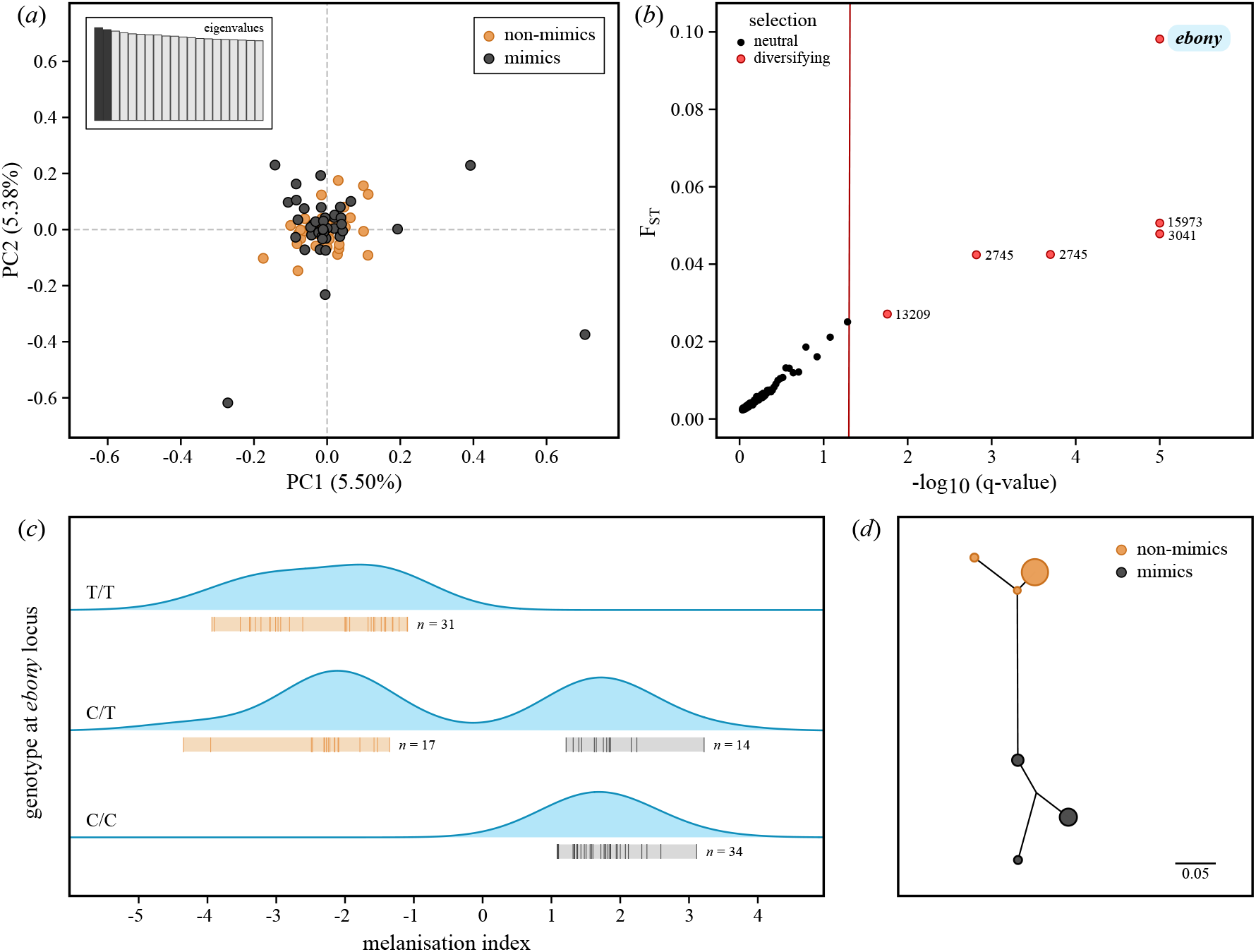
(*a*) Principal component analysis based on 83 *Zelandoperla* individuals and 7,348 neutral SNPs showing no evidence of population substructure or differentiation between mimics and non-mimics. (*b*) Results of the BayeScan outlier analysis based on 194,773 SNPs. The threshold for identifying outlier loci (q = 0.05) is indicated with a red line, and significant outliers associated with mimicry are labelled according to the identified gene, scaffold, or contig. Loci with a posterior probability of 1 were ascribed a -log_10_(q-value) of 5. (*c*) Distribution of heterozygous and homozygous genotypes at the outlier SNP located within *ebony*. (*d*) Phylogenetic relationships among mimics and non-mimics based on a 265-bp fragment of *ebony* surrounding the outlier SNP, with node size proportional to sample size.

### (d) Outlier detection and characterisation

BayeScan identified 6 SNPs putatively under diversifying selection (α > 0) among melanic and non-melanic stoneflies (Fig. 3*b*; Supplementary Table 1). Of these, only one outlier SNP (contig 10,873, position 23,588) could be associated with a known gene, occurring in a region of the reference genome orthologous to the *Drosphila* gene *ebony* (FlyBase ID: FBpp0083505). All additional outlier SNPs identified by BayeScan occurred within uncharacterised or non-coding regions of the reference genome. Based on the involvement of a highly significant outlier SNP (*ebony*) in the melanin biosynthesis pathway of insects (Wittkopp *et al*., 2002), we considered *ebony* to be a very strong candidate gene for explaining the melanic polymorphism in *Zelandoperla*. Sanger sequencing of a 265-bp fragment of *ebony* confirmed that homozygosity for the major allele (C) at the outlier SNP is restricted to mimics and homozygosity for the minor allele (T) is restricted to non-mimics, with heterozygous genotypes occurring across both groups. Analysis of surrounding genetic variation recovered five distinct haplotypes and phylogenetic analysis of these haplotypes (excluding heterozygotes) yielded two well-supported clades differentiated by 8 SNPs and a 7-bp indel (Fig. 3*d*), corresponding to mimetic versus non-mimetic phenotypes.

## Discussion

Our study highlights a rare case of *intra*specific colour-polymorphism underpinning Batesian mimicry in the wild (Mallet & Joron, 1999), with melanic individuals of palatable *Zelandoperla* closely resembling the aposematic, hydrogen cyanide-producing *Austroperla*. Crucially, genomic comparisons of mimic versus non-mimic *Zelandoperla* colour morphs detected no genome-wide divergence between them, confirming that they are conspecific. However, we detected strong divergence between these morphs at a single outlier locus (*ebony*) that has a well-documented role in insect melanism. Despite the apparent complexity of ‘warning’ colouration signals in this mimicry system (Fig 2), our findings suggest that this striking polymorphism has a relatively straightforward Mendelian genetic basis.

The gene *ebony* is well known for its key role in insect melanism, encoding for the enzyme N-β-alanyldopamine (NBAD) synthetase that converts dopamine to NBAD, which inhibits the formation of melanin, resulting in yellow pigmentation (Koch *et al*., 1998; Wittkopp *et al*., 2003). Loss of function of *ebony* thus results in darkening of the cuticle (Wittkopp *et al*., 2002; Futahashi *et al*., 2008; Tomoyasu *et al*., 2009). While *ebony* and other genes in the melanin pathway (*black, yellow, tan, Ddc, pale* [TH], *lac2*, and *aaNAT*) were originally characterised from *Drosophila* and other holometabolous model insects (Wright, 1987; Wittkopp *et al*., 2002; Arakane *et al*., 2009; Dai *et al*., 2010), recent functional analyses of melanin genes in basal insect lineages using RNA interference (Blattodea: Lemonds *et al*., 2016; Hemiptera: Liu *et al*., 2014, 2016; Zhang *et al*., 2019) support a conserved role of the melanin pathway across diverse insects, a finding reinforced by our study of an early-diverging hexapod lineage. Indeed, our detection of *ebony* involvement in a melanic stonefly polymorphism was detected despite an unbiased genome-wide approach. Future genomic analysis should shed additional light on the mechanistic basis of mutations at this and other insect pigmentation loci.

The detection of a single large-effect locus associated with melanisation in *Zelandoperla* may seem surprising, given that apparently complex phenotypic traits are frequently attributed to polygenic architectures (Fisher, 1930). While it is possible that some of the subtle variation in melanisation observed in the study population may stem from interplay between *ebony* and other genes in the insect melanin pathway (Miyagi *et al*., 2015), we have as yet no genetic evidence for the strong involvement of multiple loci. Additional genomic analyses promise to shed light on the genetic bases of variation in levels of melanism among heterozygous individuals (Fig. 3*c*).

The distinctive aposematic ‘warning’ colouration of the noxious stonefly *Austroperla* is extremely rare in the context of Plecoptera globally (Foster *et al*., 2021), making it highly unlikely that the near-identical colouration (Fig. 2) reported here in unrelated but co-distributed *Zelandoperla* represents a chance resemblance (McLellan, 1996, 1999). Additionally, the fact that the *Austroperla* is abundant in forested streams – but rare or absent from unforested habitats (Nystrom *et al*., 2003; Townsend *et al*., 1997) – provides an opportunity to test for geographic links between mimic and model occurrence (see also Tsurui-Sato *et al*., 2019). Importantly, our distributional analyses of *Zelandoperla* phenotypic records suggest that the occurrence of melanic *Z. fenestrata* is correlated with forest habitats (Fig. 1) where *Austroperla* is abundant, further supporting the suggestion that they are indeed mimics of *Austroperla*. Future studies should expand on these results by quantifying the relative abundance of mimics and models over a range of spatial scales, and further assessing genomic and phenotypic variation across the range of the *Zelandoperla* complex. We hypothesise that variation in the relative frequencies of mimic and non-mimic phenotypes of *Zelandoperla* may be closely tied to forest cover, which itself is likely a proxy for abundance of the model *Austroperla* (Townsend *et al*., 1997; Nyström *et al*., 2003). Accordingly, future studies should further test for rapid evolutionary shifts in insect colour linked to spatiotemporal environmental variation (van’t Hof *et al*., 2016).

Our analyses of *Zelandoperla* and *Austroperla* colouration and palatability clarify that their mimetic relationship is Batesian rather than Müllerian. Indeed, mimics were found to be highly palatable to predators in controlled feeding experiments, whereas individuals of *Austroperla* were essentially inedible, consistent with the well-documented chemical defences of the latter species (Thomson, 1934; Foster *et al*., 2021). Additionally, detailed morphological analyses confirm substantial overlap in melanisation and ‘warning colouration’ of mimic and model phenotypes. Specifically, melanisation in approximately 15% of the *Z. fenestrata* complex population overlapped with the range of variation recorded in *Austroperla*, with bright yellow banding on the legs of these individuals closely matching the conspicuousness of warning signals in *Austroperla*. As chemical defences and aposematic colouration in *Austroperla* likely evolved as protection against avian predators (Foster *et al*., 2021), future studies should seek to test the ability of ecologically relevant predators to learn to avoid mimics after experiences with the unpalatable model.

In conclusion, our study provides a genetic explanation for a rare intraspecific mimicry polymorphism, representing the first case of Batesian mimicry documented in Plecoptera. This mimicry system relies on enhanced contrast of ‘warning colours’ created by regional patterning of melanin pigmentation, apparently controlled by a mutation in *ebony*, a key insect melanism gene. This system thus highlights the conserved role of *ebony* across phylogenetically divergent insect lineages, and reveals an elegant intraspecific system for potentially detecting rapid adaptive shifts in response to ecological and environmental change.

## Acknowledgements

This work was supported by Marsden contracts OO1412 and UOO2016 (Royal Society of New Zealand) awarded to J.M.W. and G.A.M., and a University of Otago Doctoral Scholarship awarded to B.J.F. Specimens were collected under DOC permit 59871-RES.

## Author contributions

B.J.F. and J.M.W. designed research. B.J.F., G.A.M., and G.C.K. generated data. B.J.F. and Y.F. analysed data. B.J.F. wrote the paper. All authors edited the paper.

## References

Arakane, Y., Lomakin, J., Beeman, R.W., Muthukrishnan, S., Gehrke, S.H., Kanost, M.R., et al. 2009. Molecular and functional analyses of amino acid decarboxylases involved in cuticle tanning in Tribolium castaneum. Journal of Biological Chemistry 284: 16584– 16594.

Aronsson, M. & Gamberale-Stille, G. 2013. Evidence of signaling benefits to contrasting internal color boundaries in warning coloration. Behavioral Ecology 24: 349–354.

Bates, H.W. 1862. Contributions to an insect fauna of the Amazon Valley. Lepidoptera: Heliconidae. Transactions of the Linnean Society of London 23: 495–456.

Coffin, D. 2015. DCRAW V. 9.26 https://www.cybercom.net/~dcoffin/dcraw/.

Dai, F.Y., Qiao, L., Tong, X.L., Cao, C., Chen, P., Chen, J., et al. 2010. Mutations of an arylalkylamine-N-acetyltransferase, Bm-iAANAT, are responsible for silkworm melanism mutant. Journal of Biological Chemistry 285: 19553–19560.

Danecek, P., Auton, A., Abecasis, G., Albers, C.A., Banks, E., DePristo, M.A., et al. 2011. The variant call format and VCFtools. Bioinformatics 27: 2156–2158.

Darriba, D., Taboada, G.L., Doallo, R. & Posada, D. 2012. jModelTest 2: More models, new heuristics and parallel computing. Nature Methods 9: 772.

Edgar, R.C. 2004. MUSCLE: multiple sequence alignment with high accuracy and high throughput. Nucleic Acids Research 32: 1792–1797.

Elshire, R.J., Glaubitz, J.C., Sun, Q., Poland, J.A., Kawamoto, K., Buckler, E.S., et al. 2011. A robust, simple genotyping-by-sequencing (GBS) approach for high diversity species. PLoS One 6: 1–9.

Evangelista, D.A., Wipfler, B., Béthoux, O., Donath, A., Fujita, M., Kohli, M.K., et al. 2019. An integrative phylogenomic approach illuminates the evolutionary history of cockroaches and termites (Blattodea). Proceedings of the Royal Society B: Biological Sciences 286: 20182076.

Ferguson, L.C., Maroja, L. & Jiggins, C.D. 2011. Convergent, modular expression of ebony and tan in the mimetic wing patterns of Heliconius butterflies. Development Genes and Evolution 221: 297–308.

Fisher, R. 1930. The Genetical Theory of Natural Selection. Clarendon Press, Oxford, UK.

Foll, M. & Gaggiotti, O. 2008. A genome-scan method to identify selected loci appropriate for both dominant and codominant markers: a Bayesian perspective. Genetics 180: 977– 993.

Foster, B.J., McCulloch, G.A. & Waters, J.M. 2021. Evidence for aposematism in a southern hemisphere stonefly family (Plecoptera: Austroperlidae). Austral Entomology 60: 267– 275.

Futahashi, R., Sato, J., Meng, Y., Okamoto, S., Daimon, T., Yamamoto, K., et al. 2008. yellow and ebony are the responsible genes for the larval color mutants of the silkworm Bombyx mori. Genetics 180: 2057–2065.

Gautier, M., Yamaguchi, J., Foucaud, J., Loiseau, A., Ausset, A., Facon, B., et al. 2018. The genomic basis of color pattern polymorphism in the harlequin ladybird. Current Biology 28: 3296-3302.e7.

Geweke, J. 1992. Evaluating the accuracy of sampling-based approaches to the calculation of posterior moments. In: Bayesian statistics (J. M. Bernardo, A. F. M. Smith, A. P. Dawid, & J. O. Berger, eds), pp. 169–193. Oxford University Press, Oxford, UK.

Goudet, J. 2005. HIERFSTAT, a package for R to compute and test hierarchical F-statistics. Molecular Ecology Notes 5: 184–186.

Hoekstra, H.E. 2006. Genetics, development and evolution of adaptive pigmentation in vertebrates. Heredity 97: 222–234.

Huerta-Cepas, J., Szklarczyk, D., Forslund, K., Cook, H., Heller, D., Walter, M.C., et al. 2016. EGGNOG 4.5: A hierarchical orthology framework with improved functional annotations for eukaryotic, prokaryotic and viral sequences. Nucleic Acids Research 44: D286–D293.

Joron, M. & Mallet, J.L.B. 1998. Diversity in mimicry: paradox or paradigm? Trends in Ecology & Evolution 13: 461–466.

Kearse, M., Moir, R., Wilson, A., Stones-Havas, S., Cheung, M., Sturrock, S., et al. 2012. Geneious Basic: an integrated and extendable desktop software platform for the organization and analysis of sequence data. Bioinformatics 28: 1647–1649.

Kikuchi, D.W. & Pfennig, D.W. 2010. High-model abundance may permit the gradual evolution of Batesian mimicry: an experimental test. Proceedings of the Royal Society B: Biological Sciences 277: 1041–1048.

Koch, P.B., Keys, D.N., Rocheleau, T., Aronstein, K., Blackburn, M., Carroll, S.B., et al. 1998. Regulation of dopa decarboxylase expression during colour pattern formation in wild-type and melanic tiger swallowtail butterflies. Development 125: 2303–2313.

Krueger, F. 2015. Trim Galore: a wrapper tool around Cutadapt and FastQC to consistently apply quality and adapter trimming to FastQ files. Available at: www.bioinformatics.babraham.ac.uk/projects/trim_galore.

Kunte, K., Zhang, W., Tenger-Trolander, A., Palmer, D.H., Martin, A., Reed, R.D., et al. 2014. doublesex is a mimicry supergene. Nature 507: 229–232.

Kuznetsova, A., Brockhoff, P.B. & Christensen, R.H.B. 2017. lmerTest package: tests in linear mixed effects models. Journal of Statistical Software 82, 1–26.

Lemonds, T.R., Liu, J. & Popadić, A. 2016. The contribution of the melanin pathway to overall body pigmentation during ontogenesis of Periplaneta americana. Insect Science 23: 513–519.

Li, H. & Durbin, R. 2009. Fast and accurate short read alignment with Burrows-Wheeler transform. Bioinformatics 25: 1754–1760.

Liu, J., Lemonds, T.R. & Popadić, A. 2014. The genetic control of aposematic black pigmentation in hemimetabolous insects: insights from Oncopeltus fasciatus. Evolution and Development 16: 270–277.

Liu, J., Lemonds, T.R., Marden, J.H. & Popadić, A. 2016. A pathway analysis of melanin patterning in a hemimetabolous insect. Genetics 203: 403–413.

Mallet, J. & Joron, M. 1999. Evolution of diversity in warning color and mimicry: polymorphisms, shifting balance, and speciation. Annual Review of Ecology and Systematics 30: 201–233.

Martin, M. 2011. Cutadapt removes adapter sequences from high-throughput sequencing reads. EMBnet Journal 17: 10–12.

McCulloch, G. 2011. Evolutionary genetics of southern stoneflies. Unpublished PhD thesis, Department of Zoology, University of Otago, Dunedin, New Zealand.

McCulloch, G.A., Foster, B.J., Dutoit, L., Harrop, T.W.R., Guhlin, J., Dearden, P.K., et al. 2021. Genomics reveals widespread ecological speciation in flightless insects. Systematic Biology 70: 863–876.

McCulloch, G.A., Foster, B.J., Dutoit, L., Ingram, T., Hay, E., Veale, A.J., et al. 2019. Ecological gradients drive insect wing loss and speciation: the role of the alpine treeline. Molecular Ecology 28: 3141–3150.

McCulloch, G.A., Wallis, G.P. & Waters, J.M. 2009. Do insects lose flight before they lose their wings? Population genetic structure in subalpine stoneflies. Molecular Ecology 18: 4073–4087.

McLellan, I.D. 1997. Austroperla cyrene Newman (Plecoptera: Austroperlidae). Journal of the Royal Society of New Zealand 27: 271–278.

McLellan, I.D. 1998. A revision of Acroperla (Plecoptera: Zelandoperlinae) and removal of species to Taraperla new genus. New Zealand Journal of Zoology 25: 185–203.

McLellan, I.D. 1999. A revision of Zelandoperla Tillyard (Plecoptera: Gripopterygidae: Zelandoperlinae). New Zealand Journal of Zoology 26: 199– 219.

Miyagi, R., Akiyama, N., Osada, N. & Takahashi, A. 2015. Complex patterns of cis-regulatory polymorphisms in ebony underlie standing pigmentation variation in Drosophila melanogaster. Molecular Ecology 24: 5829–5841.

Nadeau, N.J., Pardo-Diaz, C., Whibley, A., Supple, M.A., Saenko, S. v., Wallbank, R.W.R., et al. 2016. The gene cortex controls mimicry and crypsis in butterflies and moths. Nature 534: 106–110.

Nosil, P., Villoutreix, R., de Carvalho, C.F., Farkas, T.E., Soria-Carrasco, V., Feder, J.L., et al. 2018. Natural selection and the predictability of evolution in Timema stick insects. Science 359: 765–770.

Nyström, P., McIntosh, A.R. & Winterbourn, M.J. 2003. Top-down and bottom-up processes in grassland and forested streams. Oecologia 136: 596–608.

Plummer, M., Best, N., Cowles, K. & Vines, K. 2006. CODA: convergence diagnosis and output analysis for MCMC. R News 6: 7–11.

Protas, M.E. & Patel, N.H. 2008. Evolution of coloration patterns. Annual Review of Cell and Developmental Biology 24: 425–446.

Pum Lee, K. & Wilson, K. 2006. Melanism in a larval Lepidoptera: repeatability and heritability of a dynamic trait. Ecological Entomology 31: 196–205.

Purcell, S., Neale, B., Todd-Brown, K., Thomas, L., Ferreira, M.A.R., Bender, D., et al. 2007. PLINK: a tool set for whole-genome association and population-based linkage analyses. American Journal of Human Genetics 81: 559–575.

Rebeiz, M., Pool, J.E., Kassner, V.A., Aquadro, C.F. & Carroll, S.B. 2009. Stepwise modification of a modular enhancer underlies adaptation in a Drosophila population. Science 326: 1663–1667.

Rochette, N.C., Rivera-Colón, A.G. & Catchen, J.M. 2019. Stacks 2: analytical methods for paired-end sequencing improve RADseq-based population genomics. Molecular Ecology 28: 4737–4754.

Ruxton, G.D., Sherratt, T.N. & Speed, M.P. 2004. Avoiding Attack: The Evolutionary Ecology of Crypsis, Warning Signals and Mimicry. Oxford University Press, Oxford, UK.

Schneider, C.A., Rasband, W.S. & Eliceiri, K.W. 2012. NIH Image to ImageJ: 25 years of image analysis. Nature Methods 9: 671–675.

Stavenga, D.G., Leertouwer, H.L., Hariyama, T., de Raedt, H.A. & Wilts, B.D. 2012. Sexual dichromatism of the damselfly Calopteryx japonica caused by a melanin-chitin multilayer in the male wing veins. PLoS One 7: e49743.

Tillyard, R. 1923. The stoneflies of New Zealand (Order Perlaria), with descriptions of new genera and species. Transactions and Proceedings of the New Zealand Institute 54: 197– 217.

Tomoyasu, Y., Arakane, Y., Kramer, K.J. & Denell, R.E. 2009. Repeated co-options of exoskeleton formation during wing-to-elytron evolution in beetles. Current Biology 19: 2057–2065.

Tong, J.K., Duchêne, S., Ho, S.Y.W. & Lo, N. 2015. Comment on “Phylogenomics resolves the timing and pattern of insect evolution”. Science 349: 487.

Townsend, C.R., Arbuckle, C.J., Crowl, T.A. & Scarsbrook, M.R. 1997. The relationship between land use and physicochemistry, food resources and macroinvertebrate communities in tributaries of the Taieri River, New Zealand: a hierarchically scaled approach. Freshwater Biology 37: 177–191.

Troscianko, J. & Stevens, M. 2015. Image calibration and analysis toolbox – a free software suite for objectively measuring reflectance, colour and pattern. Methods in Ecology and Evolution 6: 1320–1331.

Tsurui-Sato, K., Sato, Y., Kato, E., Katoh, M., Kimura, R., Tatsuta, H., et al. 2019. Evidence for frequency-dependent selection maintaining polymorphism in the Batesian mimic Papilio polytes in multiple islands in the Ryukyus, Japan. Ecology and Evolution 9: 5991–6002.

Untergasser, A., Cutcutache, I., Koressaar, T., Ye, J., Faircloth, B.C., Remm, M., et al. 2012. Primer3—new capabilities and interfaces. Nucleic Acids Research 40: e115.

van’t Hof, A.E. & Saccheri, I.J. 2010. Industrial melanism in the peppered moth is not associated with genetic variation in canonical melanisation gene candidates. PLoS One 5: e10889.

van’t Hof, A.E., Campagne, P., Rigden, D.J., Yung, C.J., Lingley, J., Quail, M.A., et al. 2016. The industrial melanism mutation in British peppered moths is a transposable element. Nature 534: 102–105.

van’t Hof, A.E., Reynolds, L.A., Yung, C.J., Cook, L.M. & Saccheri, I.J. 2019. Genetic convergence of industrial melanism in three geometrid moths. Biology Letters 15: 20190582.

Villoutreix, R., de Carvalho, C.F., Soria-Carrasco, V., Lindtke, D., De-La-Mora, M., Muschick, M., et al. 2020. Large-scale mutation in the evolution of a gene complex for cryptic coloration. Science 369: 460–466.

Wickham, H. 2016. ggplot2: Elegant Graphics for Data Analysis. Springer-Verlag, New York, USA.

Williams, D.S. 1979. The feeding behaviour of New Zealand Dolomedes species (Araneae: Pisauridae). New Zealand Journal of Zoology 6: 95–105.

Wittkopp, P.J., True, J.R. & Carroll, S.B. 2002. Reciprocal functions of the Drosophila Yellow and Ebony proteins in the development and evolution of pigment patterns. Development 129: 1849–1858.

Wittkopp, P.J., Williams, B.L., Selegue, J.E. & Carroll, S.B. 2003. Drosophila pigmentation evolution: divergent genotypes underlying convergent phenotypes. Proceedings of the National Academy of Sciences 100: 1800–1813.

Wright, T.R.F. 1987. The genetics of biogenic amine metabolism, sclerotization, and melanization in Drosophila melanogaster. Advances in Genetics 24: 127–222.

Zhang, Y., Li, H., Du, J., Zhang, J., Shen, J. & Cai, W. 2019. Three melanin pathway genes, TH, yellow, and aaNAT, regulate pigmentation in the twin-spotted assassin bug, Platymeris biguttatus (Linnaeus). International Journal of Molecular Sciences 20: 2728.

Zvereva, E.L. & Kozlov, M.V. 2016. The costs and effectiveness of chemical defenses in herbivorous insects – a meta-analysis. Ecological Monographs 86: 107–124.

Zwickl, D.J. 2006. Genetic algorithm approaches for the phylogenetic analysis of large biological sequence datasets under the maximum likelihood criterion. Unpublished PhD thesis, The University of Texas at Austin.

