## Supplementary methods, figures, and tables for "*Ebony* underpins Batesian mimicry in an insect melanic polymorphism"

#### Digital photography

Photographs were taken in RAW format using a Canon 40D camera with an EF-S 60 mm f/2.8 lens under fixed magnification (0.7:1), exposure time (1/6), aperture (f/9), and ISO sensitivity (400) settings. Uniform lighting was provided by a Schott-Fostec DCR II 150 W fibre optic illuminator split through four guides, which were orientated equidistantly around the subject and diffused through a polystyrene foam cylinder 60 mm in diameter. Stoneflies were restrained on pins and photographed at a standardised distance and angle on a 18% reflectance neutral grey background alongside a matte ColorGauge Pico standardised colour target (Image Science Associates, Williamson, NY, USA). Following image capture, each individual was placed in an individually labelled 1.5 mL Eppendorf tube and preserved in absolute ethanol for possible DNA extraction after selection of a subset of individuals for genotyping. After normalising images in the Multispectral Image Calibration and Analysis (MICA) toolbox (Troscianko & Stevens, 2015), the DCRAW plugin (Coffin, 2015) of ImageJ (Schneider *et al.*, 2012) was used to convert RAW files into 48-bit TIFF files for equalisation of colour values in Adobe Photoshop CC 2021 (Adobe Systems, San Jose, CA, USA).

#### Colour measurements and selection of individuals for genotyping

RGB values were sampled from each image (3888 x 2592 pixels) using the circular selection tool in ImageJ. Mean RGB values were extracted from two different sizes of the circular selection tool depending on the region of interest (ROI) (main text Fig. 2*b*). Specifically, RGB measurements associated with body melanisation (ROIs 1, 2, and 3) were averaged across a circle 30 pixels (~240 µm) in diameter while measurements associated with warning colours in *A. cyrene* (ROIs 4 and 5) were averaged across a circle 20 pixels (~160 µm) in diameter. After conversion of RGB values to HSB colour space, an index of overall body melanisation was derived from a principal component analysis of brightness values extracted from the first three ROIs. The position of 338 individuals within the melanisation index was then ranked to select a subset of individuals for genotyping-by-sequencing. Individuals with high values in the melanisation index that lacked melanisation in areas that might detract from their resemblance to *A. cyrene* (e.g. the rectangular patches between the crossveins of forewings; McLellan, 1999) were rejected from the selection procedure to ensure that genome-wide association (GWA) analyses pertained to patterns of melanisation most likely associated with mimicry. Likewise, individuals with low values in the melanisation index that had increased melanisation in areas outside of the measured ROIs were rejected. From this, 48 of the most highly melanised and 48 of the least melanised individuals fitting these criteria ('mimics' and 'non-mimics', respectively) were selected for inclusion in the GWA study.

#### Palatability experiment

Fifteen wild-caught individuals of *D. aquaticus* were kept individually in containers for 10 days and given food (crickets) and water *ad libitum* to acclimatise them to the laboratory environment (14:10 light-dark cycle, 26 °C). Adults of *A. cyrene* and the *Z. fenestrata* complex were reared continuously throughout the duration of the experiment (ambient light cycle, 11°C) to provide suitable prey for spiders on an intermittent basis. Predators were food deprived for 4 days before being presented with experimental prey for the first time. On the day of feeding, each spider was transferred to a small circular arena (90 mm diameter x 50 mm height) at the transition to the dark cycle and given 1 hour to acclimate before being presented with a randomly allocated mimic, non-mimic, or model (*A. cyrene*). The live mass of each prey item was measured immediately before feeding trials to the nearest 0.01 mg using a micro-analytical balance (Mettler-Toledo AX205 Delta Range, Greifensee, Switzerland). Feeding was observed during the dark cycle with the aid of a dim red light. Spiders were allowed time to detect and attack prey and, once feeding was initiated, were given 1 hour to finish feeding before the discarded remains of prey were weighed for comparison with their original live mass. The assays continued, with predators undergoing 4-day periods of food deprivation between feeding trials of randomly allocated prey, until the palatability of 19 mimics, 20 non-mimics, and 14 models (*A. cyrene*) had been assessed.

### Supplementary References

Coffin, D. 2015. DCRAW V. 9.26 https://www.cybercom.net/~dcoffin/dcraw/.

McLellan, I.D. 1999. A revision of *Zelandoperla* Tillyard (Plecoptera: Gripopterygidae: Zelandoperlinae). *New Zealand Journal of Zoology* **26**: 199–219.

Schneider, C.A., Rasband, W.S. & Eliceiri, K.W. 2012. NIH Image to ImageJ: 25 years of image analysis. *Nature Methods* **9**: 671–675.

Troscianko, J. & Stevens, M. 2015. Image calibration and analysis toolbox – a free software suite for objectively measuring reflectance, colour and pattern. *Methods in Ecology and Evolution* **6**: 1320–1331.

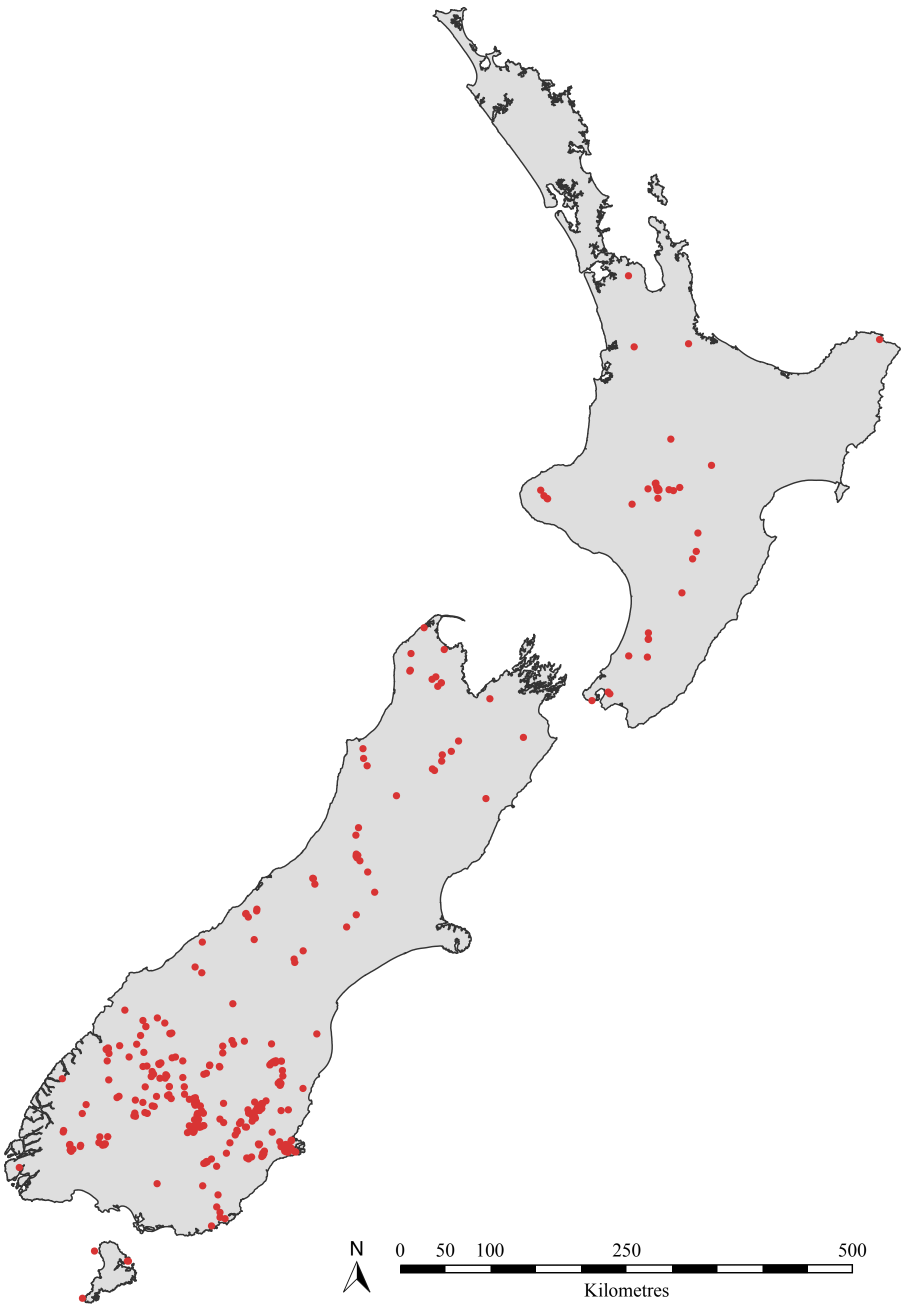


**Supplementary Figure 1.** Map showing 430 unique locations in which the colouration of 1,696 adults of the *Z. fenestrata* species complex were examined. Material was sourced from a combination of recent field collections, two private collections, and five institutional collections: Auckland War Memorial Museum, New Zealand Arthropod Collection (Auckland), Museum of New Zealand Te Papa Tongarewa (Wellington), Canterbury Museum (Christchurch), and Otago Museum (Dunedin).

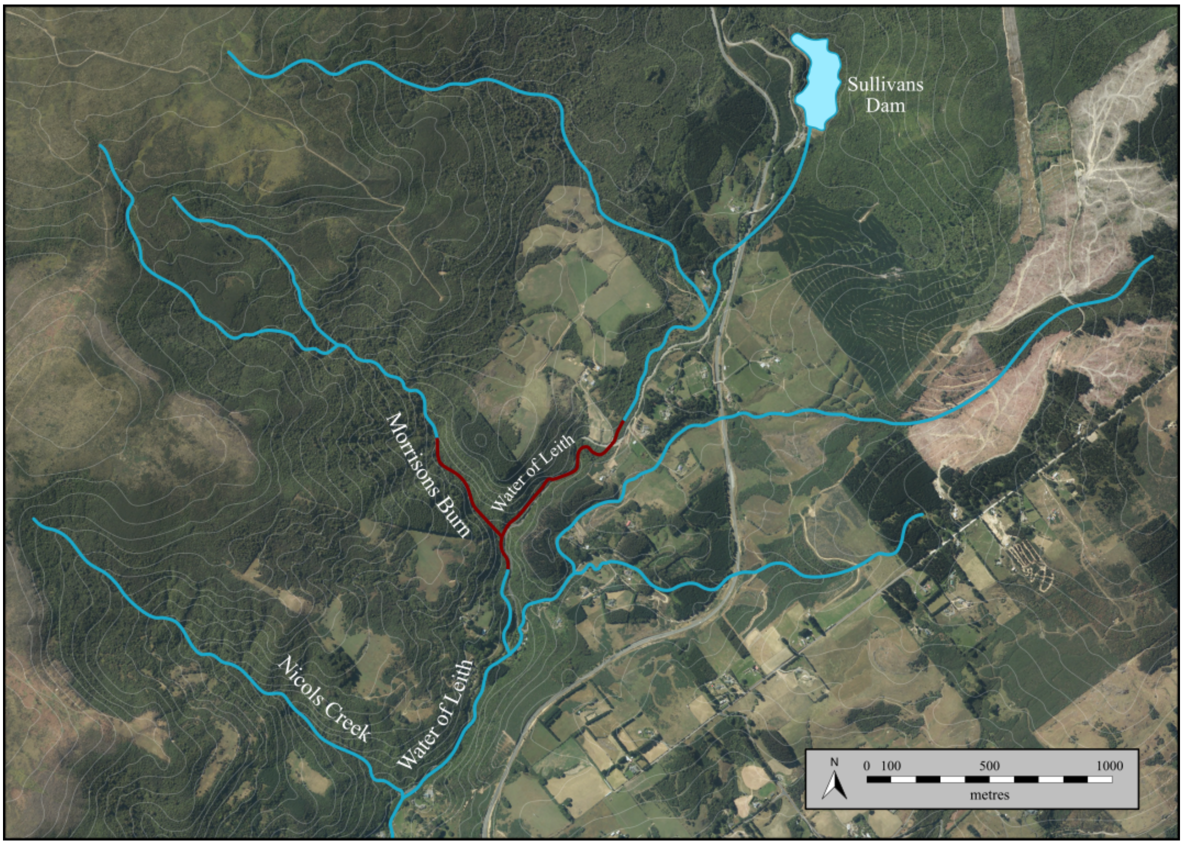


**Supplementary Figure 2.** Map of the study area in the Water of Leith catchment, Dunedin (see star in main text Fig. 1). Red segments show the areas of the Water of Leith and Morrisons Burn where nymphs of the *Z. fenestrata* species complex were collected for rearing. Contours represent 20 metre intervals in elevation (aerial photograph and elevation data sourced from the LINZ Data Service).

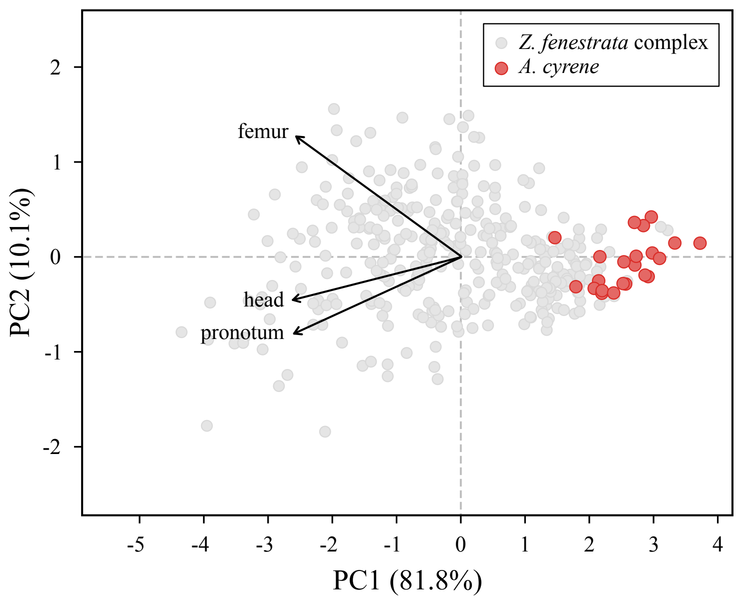


**Supplementary Figure 3**. Principal component analysis of brightness data obtained from 360 individuals (*Z. fenestrata* species complex: *n* = 338; *A. cyrene*: *n* = 22). Higher values on the first principal component correspond to decreasing brightness in three measured regions of interest (ROIs) representative of overall body melanisation. Arrows represents the direction and strength of eigenvectors for ROIs on the head, femorae, and pronotum.

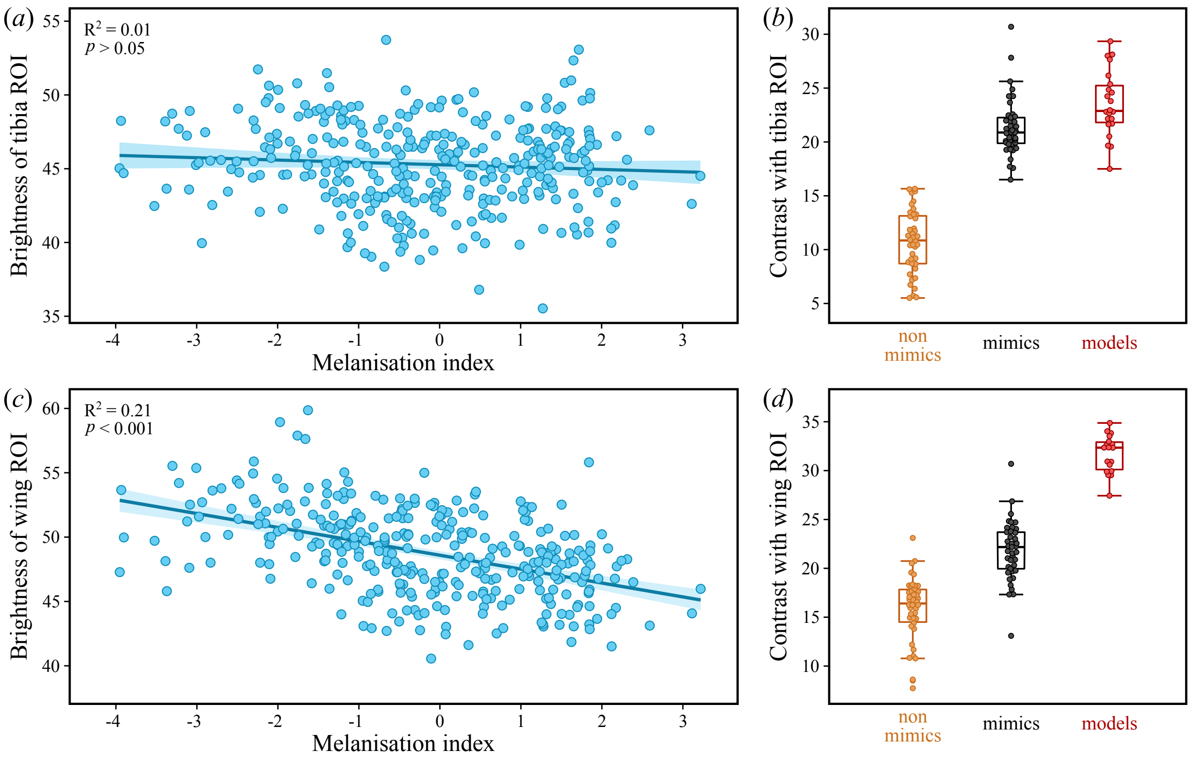


**Supplementary Figure 4**. Visual components other than body melanisation affecting the resemblance of individuals in the *Z. fenestrata* species complex to *A. cyrene*. (*a, c*) Associations between overall body melanisation and the brightness of 'warning colour ROIs' on the tibiae (*a*) and forewings (*b*) of 338 individuals in the *Z. fenestrata* species complex, evaluated using linear regression with Bonferroni adjusted *p*-values for multiple comparisons. (*b, d*) Contrast between body melanisation and colouration on the tibiae (*b*) and forewings (*d*), comparing models (*A. cyrene*) with mimics and non-mimics included in GWA analyses.

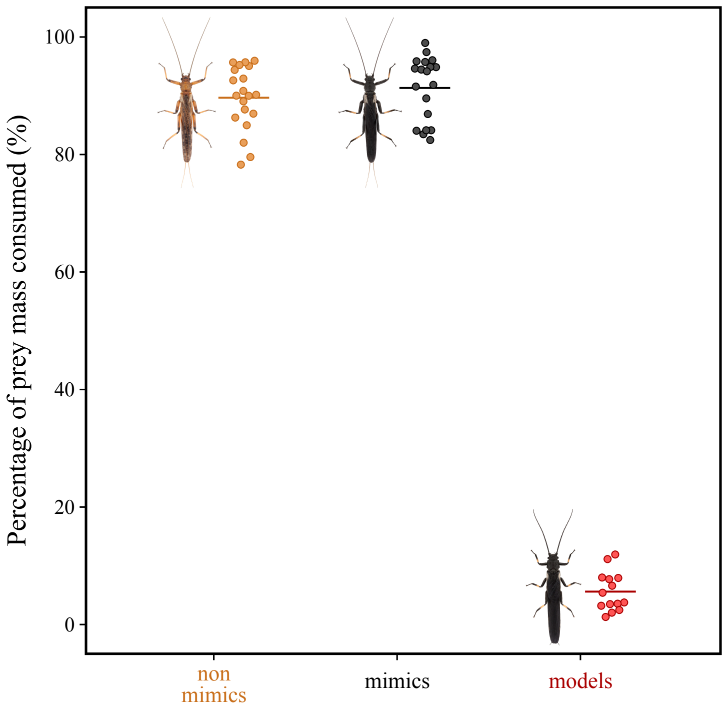


**Supplementary Figure 5**. Comparisons of palatability among mimics and non-mimics in the *Z. fenestrata* species complex and the model species, *A. cyrene*. Palatability was measured as the percentage of the mass of each prey item that was consumed by predators (*Dolomedes aquaticus*) in the absence of any visual cues. Bars represent mean values.

**Supplementary Table 1.** Outlier SNPs putatively associated with melanism identified by BayeScan.

| Contig | Contig size (kb) | SNP position (bp) | N | Alleles | MAF | FDR ($q$) | Nearest gene | Location relative to gene |
|---|---|---|---|---|---|---|---|---|
| 10,873 | 102,838 | 23,588 | 43 | C > T | 0.2558 | 0.0001 | *ebony* (SFLY2_001272) | within gene |
| 13,209 | 93,151 | 70,367 | 70 | T > C | 0.2929 | 0.0174 | unknown (SFLY2_004340) | 7 kb upstream |
| 15,973 | 19,783 | 12,538 | 56 | G > C | 0.1875 | 0.0001 | none annotated on contig | |
| 2,745 | 27,802 | 188,690 | 65 | C > A | 0.1000 | 0.0002 | unknown (SFLY2_017407) | within gene |
| 2,745 | 27,802 | 229,798 | 52 | C > T | 0.4615 | 0.0015 | unknown (SFLY2_017408) | 15 kb upstream |
| 3,041* | 417,928 | 247,562 | 62 | C > A | 0.3226 | 0.0001 | unknown (SFLY2_034908) | 5 kb upstream |

Alleles are represented as major alleles > minor alleles; N, number of individuals genotyped per SNP; MAF, minor allele frequency; FDR, false discovery rate.
*Scaffold.
